# Optimising future rhino population management strategies using insights from genetic health assessments across India

**DOI:** 10.1101/2024.08.09.607316

**Authors:** Tista Ghosh, Parikshit Kakati, Amit Sharma, Samrat Mondol

## Abstract

Various species conservation paradigms are facing enormous challenges during the ongoing Anthropocene. While the widely-used reintroduction/translocation-based approaches have supported many endangered species population recoveries, they seldom use detailed genetic information during initial planning. The Indian greater one-horned rhino typifies such assisted migration-driven species recovery, but currently facing long-term survival concerns due to their mostly small, isolated populations reaching respective carrying capacities. We assessed nation-wide rhino genetic health, identified suitable source populations and provided future translocation scenarios for all extant and proposed rhino habitats. Analyses with 504 unique rhino genotypes across all seven Indian rhino-bearing parks revealed six genetically-isolated populations with overall moderately low genetic diversity. Our results showed that Kaziranga and Manas NPs (Assam) to have the best rhino genetic health, whereas Jaldapara and Gorumara NPs (West Bengal) undergoing strong genetic erosions. Forward genetic simulations suggested that annual supplementation efforts from only few Assam rhino populations (Kaziranga NP, Orang NP and Pobitora WLS) are best suited for genetic rescue of most of the extant populations. Overall, the genetic diversity and differentiation patterns mimics the complex evolutionary history and individual recovery histories. We suggest park-specific management solutions (ranging from protection measures, grassland restoration, livestock and conflict management, regular supplementation events etc.) to ensure the species’ long-term persistence and prevent the alarming loss of grassland habitats and its associate biodiversity. We insist on utilising such genetic health indices-driven population management solutions to identify targeted mitigative measures in other species.

## 1. Introduction

The continuing anthropogenic pressures on natural systems have resulted in massive declines in species diversity and ecological services globally (Exposito-Alonso et al., 2023; Le Roux et al., 2018; Malhi et al., 2015; WWF 2020). Accordingly, species conservation paradigms have also adapted towards multifaceted management approaches to reduce the impacts of such external threats (Bolam et al., 2020; Lindsey et al., 2017; Smith et al., 2023; Sylven et al., 2012). In case of absence of natural geneflow, translocation and reintroduction-based conservation programmes have become one such widely-accepted choices for recovery of endangered, extinction-prone species (Hostetler et al., 2010; Johnson et al., 2010; O’Brien et al., 2017; Stanbridge et al., 2023; Whiting et al., 2023; Willi et al., 2022). Although the benefits of such assisted migration-based approaches are not new (Hogg et al., 2006; Johnson et al., 2010; Weeks et al., 2017; Scotts et al., 2020; Thomas et al., 2023), it is important to consider the genetic and demographic consequences of the recovered populations from founder effects to future stochastic events (Mace et al., 2010; Moodley et al., 2017; Lino et al. 2019; Stanbridge et al., 2023; Willi et al., 2022). This is critical as the global conservation paradigm often measures species recovery success only in terms of increase in population size but seldom consider integrating genetic and other demographic information (Coleman et al., 2013; Taylor et al., 2017; Whiteley et al., 2015; DeWoody et al., 2021). Large numbers of empirical and experimental research indicate direct relationship of genetic diversity with population fitness (Hufbauer et al., 2015; Kronenberger et al., 2018; Pacioni et al., 2020; DeWoody et al., 2021) but their use in existing species recovery programmes is very low (for example, ∼0.02% of species recovery programs across the US, Fitzpatrick et al., 2023), possibly due to limited information of genetic data from these species (Fitzpatrick et al., 2023). Thus, integration of population-level genetic data is essential for successful implementation of any such assisted migration-based conservation programmes (Kelly et al., 2019; White et al., 2020; Robinson et al., 2021; Rezic et al.,2022).

The greater one-horned rhinoceros (*Rhinoceros unicornis*) is one such global example of a successful translocation and effective conservation-driven (Rookmaker et al., 2016) population recovery. Currently restricted to only 11 isolated populations along the Terai-Duars-lower Brahmaputra landscape (covering Nepal, north and north-east India) the species has revived from few hundreds in late 90s to ∼3270 individuals at present (Menon, 1996; Rookmaker et al., 2016; Sharma, 2022). Given that India retains ∼81% of the species global wild population (Sharma, 2022), the conservation actions taken by India would be critical for the species’ future persistence. Currently majority of the Indian rhino populations (except Kaziranga NP, Assam; n=∼2500) are small (ranging from 40-280), isolated and reaching their respective carrying capacities (Jhala et al., 2021). Therefore, all major government-mandated conservation programmes are focused towards expanding rhino habitats through translocation-driven population recovery plan (National Conservation Strategy for the Indian One-horned rhinoceros (*Rhinoceros unicornis*), Government of India, Ministry of Environment Forest and Climate Change, 2021). As all the extant rhino-bearing areas have recovered from small founder populations (potentially lower Ne) and preliminary information indicate genetic isolation (at state level based on mitogenome data by Ghosh et al., 2022; at local population level for West Bengal by Das et al., 2015, for details see Table S1), detailed assessment of genetic health of these populations would be critical towards future adaptive management plans. Earlier researches have provided detailed insights into the evolutionary history of Indian rhinos (Ghosh et al., 2022) and baseline information on local rhino genetic diversity (Borthakur et al., 2016; Das et al., 2015; Zschokke et al., 2011), but these data cannot be effectively used to propose recovery plans.

Considering this background, our study focuses on investigating comprehensive genetic health assessments of all Indian rhino populations and use the information to generate targeted management suggestions and identify healthy source populations for proposed new habitats. More specifically, the objectives are to (i) estimate the extent of genetic structuring in Indian rhinos; (ii) identify the genetically vulnerable populations (based on multiple indices) and (iii) provide best possible scenarios for translocations/reintroduction to maintain high genetic variability in extant as well as proposed new rhino habitats. We addressed these questions using a multidisciplinary approach involving integrated Bayesian statistics, genetic diversity indices and forward genetic simulations on 12 STR loci data from 504 individual rhinos collected across its distribution in India.

## 2. Materials and methods

### 2.1 Permission and ethical statement

All the authorizations related to biological sampling (both tissue and dung) for this study was permitted by Ministry of Environment, Forests and Climate Change (MoEF&CC), Government of India (Letter No. 4-22/2015/WL). Permissions for field sampling was provided independently from each state forest departments of Assam (Letter No. A/GWL/RhoDIS/2017/913, 3653/ WL/2W-525/2018, WL/FE.15/22), West Bengal (Letter No. 3967/WI/2W-525/2018) and Uttar Pradesh (Letter No. 1978/23-2-12 (G)). No ethical permissions were required for the tissue samples as they were collected from naturally dead rhinos by the veterinary officers of the respective forest departments during regular post-mortem procedures.

### 2.2 Study area

The study was conducted covering all seven Indian rhino bearing areas spread across three states: Uttar Pradesh (Dudhwa NP-westernmost range), West Bengal (Gorumara and Jaldapara NPs) and Assam (Manas NP, Pobitora WLS, Orang NP and Kaziranga NP (easternmost distribution (Figure 1)). All the parks have experienced different kinds and magnitudes of anthropogenic pressures in last 200-300 years (Rookmaker et al., 2016). Some of these populations (for example, Dudhwa and Manas NPs) were locally extinct and were recovered through reintroduction efforts to current population sizes of 40 and 48, respectively (Sharma, 2022; Talukdar et al., 2021). Kaziranga NP retains the largest numbers of wild one-horned rhinoceros (Sharma, 2022) whereas the other populations host varied population sizes (40-∼300 individuals) with different recovery histories. Details of each park with respect to its recovery history, density, grassland area and connectivity status (isolated or connected) are given in <u>Table S2</u>. Such unique and varied histories of these extant populations provide an excellent opportunity to investigate the genetic outcomes resulting from a myriad of anthropogenic interventions.

**Figure 1:**
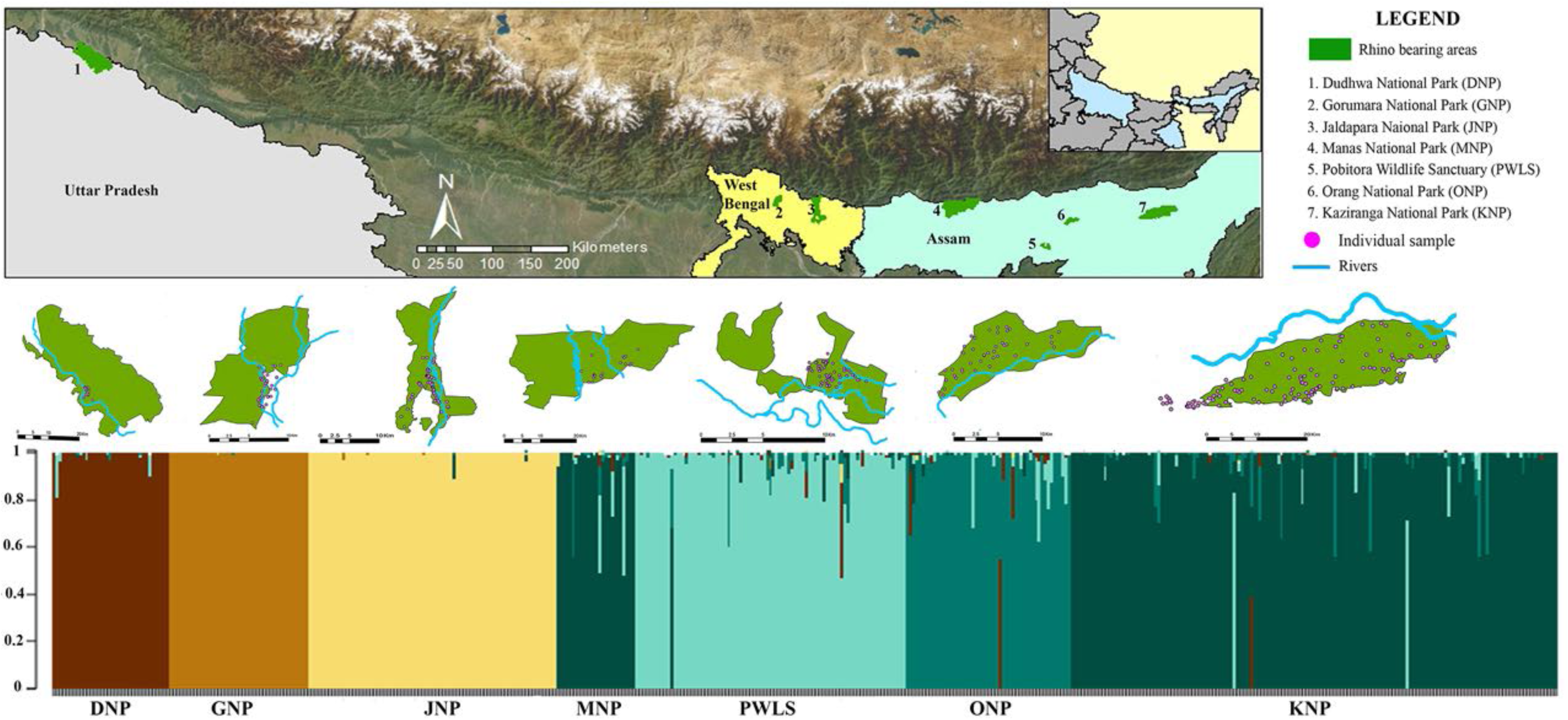
Map of the study area representing spatial distribution of 504 identified individuals and their respective genetic cluster. The top plate shows the position of the rhino-bearing parks across three Indian states (Uttar Pradesh, West Bengal and Assam). The individual location outside the boundaries of Gorumara NP and Kaziranga NP represents the population residing in the recent extended areas of these parks. The lower plate shows the identified genetic clusters i.e. K=6.

### 2.3 Sample collection

To assess rhino genetic health, we ensured fine-scale sampling coverage across their Indian distribution. We collected a total of 901 samples (153 tissues and 748 fresh dungs) between 2017-2020 (Table 1). The sampling effort was decided based on local population variables (population size and density, recovery history, grassland area, rhino distribution information from local forest staff etc.) to capture the inherent genetic gradient caused by heterogeneous species distributions and social organisations (Schwartz & McKelvey 2009; Segelbacher et al., 2010). Since rhino use communal latrines (middens), we selected the topmost fresh dung bolus of pre-identified middens during foot/vehicle surveys and swabbed (in duplicate) the top layer with sterile swabs (Himedia, Mumbai, India) to avoid cross-contamination. Additionally, tissue samples of naturally dead rhinos were provided by respective forest departments as part of the RhoDIS-India protocol. All geo-tagged samples were stored in room temperature up to a maximum period of one week (during field surveys) before they were shipped to laboratory and stored in -20°C freezer till downstream processing.

**Table 1:** Detail of sampling effort, individual identification and genetic diversity/health indices for each park.

| State | Population | Sample details |  | Genotyping error rates |  |  |  | Genetic diversity indices |  |  | Genetic health indices |  |  |  |
| --- | --- | --- | --- | --- | --- | --- | --- | --- | --- | --- | --- | --- | --- | --- |
|  |  | Sample collected | Individuals identified | AS | ADO | FA | NA | He | Ho | Fis | GD | C | Fit | Health Condition |
| Uttar Pradesh | DNP | 94 | 39 | 92 | 0.07 | 0.03 | 0.03 | 0.43 | 0.41 | 0.03 | + | + | 0.24 | Moderate |
| West Bengal | GNP | 112 | 47 | 89 | 0.07 | 0.02 | 0.03 | 0.34 | 0.3 | 0.07 | - | - | 0.45 | Poor |
|  | JNP | 121 | 83 | 89 | 0.06 | 0.03 | 0.05 | 0.41 | 0.36 | 0.08 | + | - | 0.25 | Poor |
| Assam | MNP | 78 | 26 | 82 | 0.15 | 0.02 | 0.1 | 0.52 | 0.34 | 0.29 | - | + | 0.05 | Good |
|  | PWLS | 216 | 91 | 78 | 0.16 | 0.02 | 0.14 | 0.46 | 0.24 | 0.42 | + | + | 0.11 | Moderate |
|  | ONP | 80 | 55 | 85 | 0.06 | 0.02 | 0.07 | 0.53 | 0.4 | 0.2 | - | - | 0.08 | Moderate |
|  | KNP | 200 | 163 | 92 | 0.04 | 0.02 | 0.08 | 0.59 | 0.48 | 0.15 | - | + | 0.05 | Good |

### 2.4 DNA extraction, PCR amplification and data quality controls

DNA was extracted from all field-collected tissue and dung swabs following earlier standardized protocols (Ghosh et al., 2022). In brief, samples were digested overnight with combination of ATL and Proteinase K (20 mg/ml) at 56 °C, followed by modified QIAamp DNA Tissue Kit (QIAGEN Inc., Hilden, Germany) protocol (Biswas et al., 2019). DNA was eluted twice in 100 μl preheated (70°C) 1X TE buffer and stored in -20°C freezer. Extraction negative was used for each set of experiment (n = 23) to check contamination.

Unique rhinos were identified using an existing forensic panel of 14 microsatellites earlier described (Ghosh et al., 2021). The PCR was conducted in 10μL reactions containing 4μL of 2X Qiagen multiplex PCR buffer mix (QIAGEN Inc., Hilden, Germany), 0.25μM of primer mix, 4μM BSA (4 mg/mL) and 2μL of rhino DNA. PCR conditions included an initial denaturation (95°C for 15 min); 40 cycles of denaturation (95°C for 30 s), annealing (Ta °C for 40 s) and extension (72°C for 40 s); followed by a final extension (72°C for 30 min). During each reaction set, PCR and extraction negatives were included to monitor contamination. As per the RhoDIS-India protocol (Ghosh et al., 2021), an existing reference rhino DNA sample was used in each reaction plate to ensure data uniformity. Amplified products were genotyped using HiDi formamide (Applied Biosystems, California, United States) and LIZ 500 size standard (Applied Biosystems, California, United States) in an ABI 3500XL Genetic Analyser (Applied Biosystems, California, United States). All the samples were genotyped thrice independently and manually scored in GENEMARKER (Softgenetics Inc., Pennsylvania, United States).

Data accuracy was assured through a modified ‘Quality index’ approach where a sample-wise threshold of 0.66 per locus and 0.75 mean across all the loci was implemented (Ghosh et al., 2021, Modi et al., 2021). Genetic recaptures were removed using program CERVUS (Kalinowski et al., 2007), followed by calculation of probability of identity for siblings (PID(sibs)) and unbiased (PID(unbiased)) through program GIMLET (Valiere, 2002). Amplification success, allelic dropout (ADO) and false allele (FA) rates were manually quantified (Broquet & Petit 2004; Pompanon et al., 2005). Null allele (NA) was estimated using EM algorithm implemented in program FreeNA (Chapuis & Estoup 2007). Departure from Hardy Weinberg equilibrium and linkage disequilibrium was checked using ARLEQUIN 3.1 (Excoffier et al., 2005).

### 2.5 Population structure and genetic diversity

Considering the evolutionary history (Ghosh et al., 2022), small founder gene pool and varied recovery patterns of the extant rhino populations (Rookmaker et al., 2016), we used an integrated clustering approach to understand the patterns of genetic structuring (Basto et al., 2016; De et al., 2021; Hobbs et al., 2011; Paul et al., 2023a). We implemented Bayesian clustering methods based on admixture (programs STRUCTURE (Pritchard et al., 2000) and TESS (Chen et al., 2007)) and non-admixture (program GENELAND (Guillot et al., 2005)) models to identify genetic partitions even with low level of differentiations (Basto et al., 2016; Gaggiotti et al., 2009; Schwartz et al., 2009). Some of these models are known to produce under or overestimated K values (Basto et al., 2016; Schwartz et al., 2009; Vergara et al., 2015) due to inherent prior assumptions of population genetic models. Therefore, we used a multivariate analysis implemented in program sPCA (Jombart, 2008) that accounts for spatial autocorrelation issues related with Bayesian approaches (Jombart, 2008; Vergara et al., 2015). We used a consensus approach collating all the results to infer the K value (Basto et al., 2016; De et al., 2021; Paul et al., 2023a).

Following run parameters were used in the analyses: (a) STRUCTURE v2.3 was run including with and without Locprior assumptions (Pritchard et al., 2000) for 10 independent runs of each K value (1 to 12) with 4.5 X 10^5^ iterations, 5X10^4^ burn-in and correlated allele frequencies. The optimal K was selected using the L(K) statistics (Pritchard et al., 2000) after model averaging 12 replicates using CLUMPAK (Kopelman et al., 2015); (b) TESS v2.3.1 run parameters included 10 replicate runs of 25000 sweeps after 1200 burn-in to test the most likely number of clusters (KMAX=2 to 12). Model averaging was done using CLUMPAK (Kopelman et al., 2015) and KMAX was selected based on the deviance information criterion (DIC) (Durand et al., 2009). Results of both the programme were visualised with STRUCTURE PLOT v2.0 (Ramasamy et al., 2014); (c) for GENELAND v4.0.8 simulations of 2X10^5^ iterations and 200 thinning were used without uncertainty on spatial coordinates for K=1 to 10 (with 10 replicates for each K) employing correction for null alleles. The optimal K was decided based on the highest mean posterior density across all iterations followed by generation of individuals assignment map (Guillot et al., 2005); and (d) sPCA runs were carried to assess the significance of global spatial structure between population pairs showing contradictory results in Bayesian approaches (Jombart 2008). This was done using nearest neighbour as connection network and Monte Carlo test with 9999 iterations to get Moran’s I index.

Once genetic subpopulations were established, we estimated pairwise Fst among the clusters and conducted hierarchical analysis of molecular variance (AMOVA) using ARLEQUIN 3.1 (Excoffier et al., 2005) with 10000 permutations and testing for significance at α=0.05. Finally, we calculated basic indices of genetic diversity and inbreeding coefficient (Fis) for all the identified subpopulations using Arlequin (He, Ho) and Fstat (Fis) (Excoffier et al., 2005).

### 2.6 Assessing genetic health of Indian rhino populations

Given the small sizes of most of the extant rhino populations (except Kaziranga NP) and their varied demographic histories, it is critical to assess park-specific genetic health to develop necessary mitigation strategies. We used three summary statistics-based indices focused on inheritance of genetic diversity, allelic richness and genetic erosion, respectively, to assess genetic health of all rhino populations. Overall, the combination of these indices provides information on patterns of diversity loss and therefore help identifying genetically vulnerable population/s (De et al., 2021; Kolipakam et al., 2019; Mannise et al., 2017).

The first index used is mean genetic distance (GD) which measures the maintenance of overall Nei genetic diversity, where a positive value indicates less contribution to the overall gene pool (Alvarez et al., 2009; Caballero, & Toro, 2002; Mannise et al., 2017). The second index represents population-wise genetic uniqueness conceptualized based on Hulbert’s rarefacted allelic richness (C) (Petit et al., 1998), where a positive value indicates more contribution toward aggregated allelic diversity (or high allelic richness) (Alvarez et al., 2009; Kolipakam et al., 2019; Petit et al., 1998). These indices were calculated using MolKin (Gutíerrez et al., 2005) with the default settings. The third index, population specific Fst (Fit), is an indicator of genetic erosion (Foll, & Gaggoitti, 2006), where the values range from 0 (no genetic erosion) to 1 (complete erosion). The population-specific Fit values were estimated using GESTE with default settings for zero factor analysis (Foll, & Gaggiotti, 2006).

Finally, all the three indices were compiled to assess genetic health of all seven Indian rhino populations. The expectations from a genetically healthy population would be represented by a negative GD, positive C and lower Fit values (Alvarez et al., 2009; Foll et al., 2006; Gutíerrez et al., 2005).

### 2.7 Genetic data simulation

As all future rhino population conservation/management plans are translocation/ reintroduction-driven (National Conservation Strategy for the Indian One-horned rhinoceros (*Rhinoceros unicornis*), Government of India, Ministry of Environment Forest and Climate Change, 2021), it is critical to consider genetic information at the planning stage (Kelly et al., 2019; Rezic et al.,2022; Robinson et al., 2021; White et al., 2020). We used VORTEX 9.93 (Lacy 1993) to simulate the future trends in genetic diversity by modelling different scenarios of genetic supplementation, where various demographic parameters (such as birth rate, death rate, carrying capacity etc.) were used from earlier studies (Jhala et al., 2021; Subedi et al., 2017) along with the genetic information generated in this study. Before the simulations were conducted certain population-specific considerations were implemented to ensure best possible outputs. For example, the reintroduced populations (Manas NP and Dudhwa NP) were not considered as source for any simulation analysis as mixed genetic signatures would be expected in them (based on their source population histories). Further, during data inputs any population with low genetic health index (than the recipient one) was not considered as a source. The rhino translocations scenarios were modelled with various combination of potential sources (one to four populations), translocation frequency (annually or alternative year), supplementation time (10, 25 years) and male-female ratio (1:1 and 1:2) during translocation events (Table S4). Finally, we have also performed simulations for two future rhino reintroduction plans (in Valmiki NP, Bihar and a new population in Uttar Pradesh) to identify best sources and scenarios. In this case, we modelled populations with constant carrying capacity of 100 individuals (as suggested in Jhala et al., 2021) and different founder sizes (5-20 individuals). Overall, 1000 iterations were run for 100 years in each scenario (n=378, Table S4).

## 3. Results

### 3.1 Rhino genetic diversity and population structure

A panel of 12 loci (out of 14) was finalised for individual identification and downstream analysis as locus SR281 and 12F continuously provided inconsistent data (low amplification success and unstable peak characters resulting in <40% samples producing data for analyses). Out of 901 collected samples, 504 unique individuals were identified after removal of 265 poor quality samples and 132 recaptures (details in <u>Table S3</u>). Overall, we observed a high average amplification success (87%) and low genotyping error rates (ADO-0.09, FA-0.02, NA-0.07) across all rhino genotype data. The cumulative PID(sibs) and PID(unbiased) values were calculated as 4.6 X 10^-4^ and 2.7 X 10^-8^, respectively, indicating strong statistical support for unambiguous rhino individual identification (see <u>Table S3 for details)</u>.

For rhino population structure assessment, results varied across different Bayesian approaches (K=4-7 based on spatial and non-spatial analyses) with consensus of four distinct clusters corresponding to Gorumara NP, Jaldapara NP (West Bengal), Kaziranga NP and Pobitora WLS (Assam) with remaining parks (Dudhwa NP, Manas NP and Orang NP) showing different clustering (Table S3). The major difference between the admixture-based non-spatial (STRUCTURE, K=4) and the spatial (STRUCTURE-Locprior and TESS, K=5) was the identification of Dudhwa NP as a distinct genetic cluster. The non-admixed GENELAND analyses showed K=7 corresponding to the individual sampling locations (Figure S1). The pairwise sPCA approach indicated true genetic clusters between Dudhwa NP-Pobitora WLS (significant global pattern, p=0.001), Manas NP-Pobitora WLS (p=0.001), Orang NP-Kaziranga NP (p=0.001) and Orang NP-Pobitora WLS (p=0.001), but did not support the same for Manas NP-Kaziranga NP (p=0.19). Combined together, the results suggest the presence of six rhino genetic sub-populations (K=6): Dudhwa NP (Uttar Pradesh), Gorumara NP and Jaldapara NP (West Bengal), Orang NP, Pobitora WLS and combined Kaziranga NP-Manas NP (Assam) (Figure 1). The AMOVA results corroborate this hierarchical structure pattern in the data where 88% of the variations were found ‘within population’, thereby indicating sub-structuring. The pairwise Fst values ranged between 0.03-0.29 among these six genetic clusters (Table 2).

**Table 2:** Pairwise Fst comparisons (p<0.05) among the identified genetic sub-populations.

| **Genetic sub-populations** | **Dudhwa** | **Gorumara** | **Jaldapara** | **Manas + Kaziranga** | **Pobitora** | **Orang** |
| --- | --- | --- | --- | --- | --- | --- |
| Dudhwa NP | 0 |  |  |  |  |  |
| Gorumara NP | 0.29 | 0 |  |  |  |  |
| Jaldapara NP | 0.13 | 0.23 | 0 |  |  |  |
| Manas NP + Kaziranga NP | 0.11 | 0.21 | 0.14 | 0 |  |  |
| Pobitora WLS | 0.06 | 0.27 | 0.16 | 0.04 | 0 |  |
| Orang NP | 0.09 | 0.23 | 0.15 | 0.04 | 0.03 | 0 |

Overall, the Indian one-horned rhinos showed moderately low genetic variations (all data combined) with low numbers of alleles (3.75), and moderate observed heterozygosity (Ho: 0.38) (Table 1). However, at population-level we observed variations in certain population-specific parameters (See Table 1 for park-wise details). For example, the number of alleles ranged between 2.5±0.96 (Gorumara NP) to 3.58 ±1.50 (Kaziranga NP), whereas the inbreeding coefficient values (Fis) varied drastically (Fis_(Dudhwa NP)_-0.03 to Fis_(Pobitora WLS)_-0.42). Comparison at state level indicate Assam has the highest genetic variation, followed by Uttar Pradesh and West Bengal. We did not observe any deviations from Hardy-Weinberg Equilibrium for any of the loci.

### 3.3 Assessing genetic health of Indian rhino populations

Considering the cumulative outputs of the three genetic indices (GD, C and Fit) we categorised all the wild rhino-bearing areas into three groups of populations with Good, Moderate and Poor genetic health. Kaziranga and Manas NPs (Assam) showed the best genetic indices (–ve GD, +ve C and lowest Fit value (0.05)) and therefore can be considered as the best rhino population in terms of genetic health. Further, Orang NP, Pobitora WLS (Assam) and Dudhwa NP (Uttar Pradesh) rhinos showed intermediate genetic health with moderate Fit values (0.1-0.2) but loss of diversity in either of the other two indices (i.e. +GD or -C). Finally, the two population from West Bengal (Jaldapara and Gorumara NPs) showed strong signatures of genetic erosions (Fit values ≤0.25) and signs of diversity loss. Overall, the health matrix (in ascending Fit order as shown in Table 3) indicates that both West Bengal rhino populations are genetically vulnerable and require immediate conservation attention.

### 3.4 Genetic data simulation

Simulation analyses to assess future genetic diversity trends under various population supplementation scenarios revealed certain common outcomes: (i) Few populations (Kaziranga NP, Orang NP, Pobitora WLS) are the best sources available for genetic rescue in majority of the simulation outcomes; (ii) annual supplementation plans leads to increase in genetic diversity than the alternative one; and (iii) translocating various male: female ratio and different supplementation tenures (10 vs 25 years) did not show any impacts in genetic diversities.

The genetic diversity projections for Jaldapara and Gorumara NPs (poor genetic health) produced similar outcomes for either single or multiple (mentioned above) source individuals with Kaziranga NP and Orang NP showing best results. For Orang NP and Pobitora WLS (moderately healthy populations) such long-term managed translocation-driven plans did not exhibit any differences in genetic diversity. So, we modelled metapopulation scenarios based on available information (Orang-Kaziranga:- Jhala et al., 2021; Pobitora-Orang:- Talukdar et al., 2007; and Pobitora-Orang-Kaziranga:-Talukdar et al., 2000). The results showed substantial increase in genetic diversity at Orang NP but not in Pobitora WLS (Figure 2).

**Figure 2:**
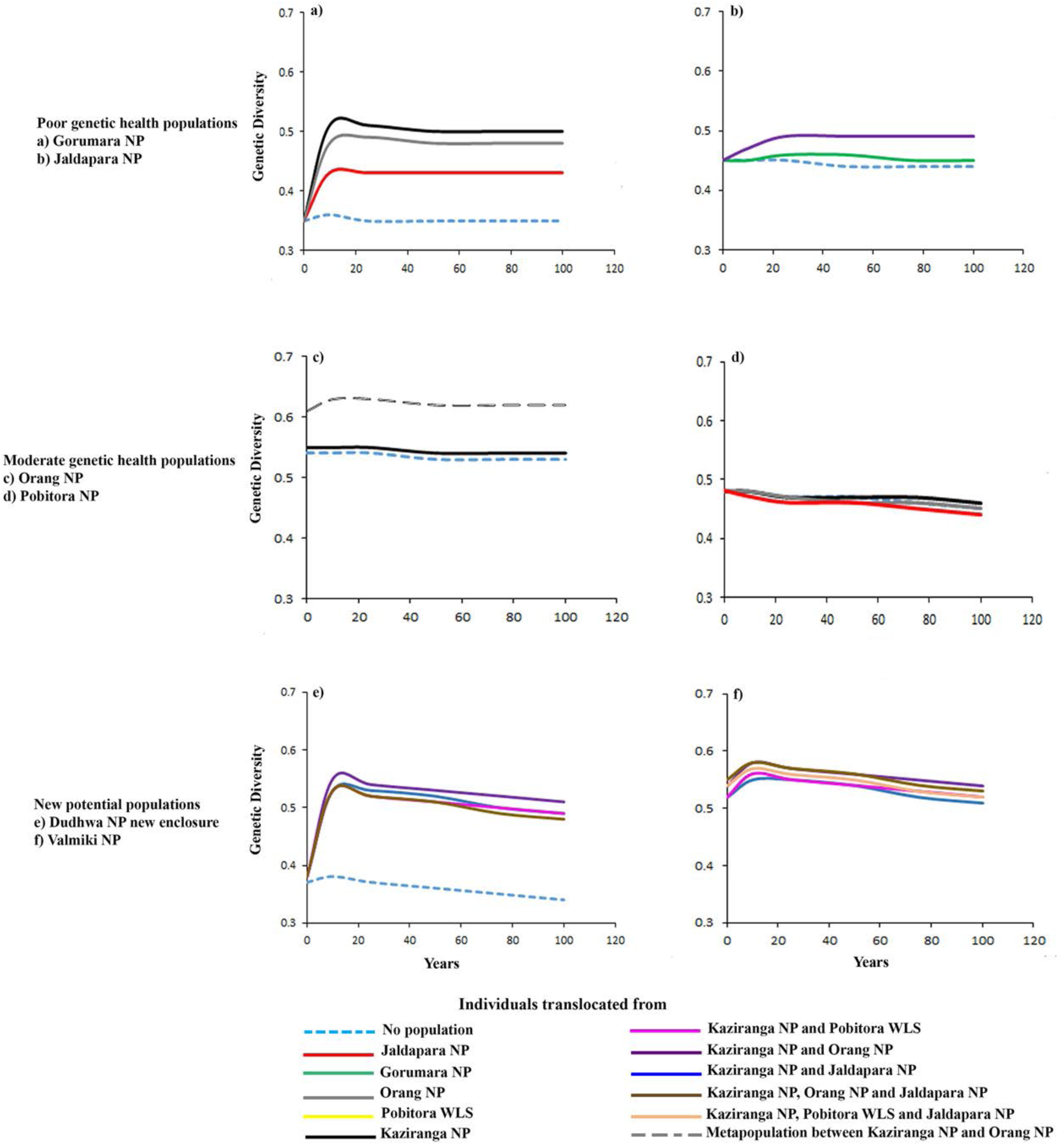
Result of simulated supplementation scenarios to assess the change in genetic diversity for populations with poor (a,b) and moderate genetic (c,d) health rank along with future potential habitats (e,f). Here we are presenting the highlights of simulation analysis for ease of data interpretation, the detailed description for all the scenarios is provided in Supplementary table 4.

Population simulations for the rhino reintroduction in new habitats (Valmiki NP, Bihar and new population in Uttar Pradesh) indicate that multiple source populations (2-4 sources, see Table S4) would result in higher genetic diversity. A long-term viable population can be maintained through introducing at least 10 genetically variable founder individuals followed by annual supplementation from varied sources (Kaziranga, Orang, Pobitora and Jaldapara-arranged in descending order of source population rank).

## 4. Discussion

The extant fragmented and isolated wild one-horned rhinoceros populations across the Indian subcontinent have experienced complex evolutionary processes to reach its current status. Fossil records indicate that the species distribution got restricted to the grassland-dominated regions of the Terai-Arc and Dooars during the Holocene climatic optimum period (Patnaik 2016). Recent genetic assessments also suggested three ‘Evolutionary Significant Units (ESUs)’ within the extant Indian rhino populations corresponding to the states of Assam, West Bengal and Uttar Pradesh, respectively, with strong genetic isolations among them (Ghosh et al., 2022). However, the nuclear marker-based assessments in this study indicated more fragmented populations across India, where our data (representing ∼15% of the total rhino population in India) showed six rhino subpopulations (Dudhwa NP, Gorumara NP, Jaldapara NP, Pobitora WLS, Kaziranga NP and Orang NP) which can possibly be explained by their recent recovery history. Such patterns of strong genetic structure mediated by both evolutionary and contemporary history has been found in other rhino species as well (for example, divergence between northern and southern white rhino, Moodley et al., 2018; Black rhino, Moodley et al., 2017; Sumatran rhino, Brandt et al., 2018; Javan rhino, Margaryan et al., 2020). Within Assam the reintroduced population of Manas NP mimics the genetic signatures of Kaziranga NP whereas the founder members were also brought from Pobitora WLS. However, monitoring records suggest higher death counts of the Pobitora individuals and subsequent translocation of more mother-calf pair from Kaziranga NP (Talukdar et al., 2020) leading to such similarity. Further, Orang NP showed a rather weak population structure (incipient structure with low Fst of 0.04) with Kaziranga NP and Pobitora WLS, possibly driven by occasional rhino immigration events from both populations (Talukdar 2000; Talukdar et al., 2007). Despite being geographically closer than any of the Assam populations, the West Bengal rhino-bearing parks (Jaldapara and Gorumara NPs) showed strong genetic differentiation (Fst 0.23). This pattern corroborated with earlier results from the same landscape (Das et al., 2015). Such outcome can possibly be a result of long genetic isolation between them since 1800s when tea plantation by British regime initiated, leading to habitat encroachment and initiation of sport/bounty hunting practices (Bist 1994; Rookmaker et al., 2016). Interestingly, Jaldapara showed lower genetic differentiation with the Assam populations (Fst ranges 0.14-0.16) than Gorumara, possibly due to historical inter-state habitat connectivity between these areas through Sankosh–Rydak–Manas landscape till 1960s (Bist 1994). Further, genetic distinction of the Dudhwa rhinos can be explained by their reintroduction history, where the first three generations were sired by one dominant male of Pobitora WLS and four females from Chitwan NP, Nepal (Talukdar et al., 2012) (leading to strong differentiation due to genetic inputs from Nepal individuals, a pattern also observed by Ghosh et al., 2022 based on mitochondrial genome). Among the Indian populations Dudhwa rhinos were found to be genetically much closer to Pobitora as compared to others (Table 2).

Overall, the genetic diversity of Indian rhinos (n=504, He= 0.56, Ho=0.38) is moderate in comparison to African (White rhino: n= 232, He= 0.62, Ho=0.47 (Moodley et al., 2018); Black rhino: n=216, He=0.61, Ho= 0.53 (Moodley et al., 2017)) and Sumatran (n=13, He=0.5, Ho=0.28 (Brandt et al., 2018)) species. Similar patterns were also reported for rhino mitogenome diversity (Ghosh et al., 2022). Earlier studies reported similar indices of genetic diversity (n=9, He= 0.45 and Ho= 0.43) from GOH rhino population of Nepal. However, it is important to realise that the basic summary statistics used for such comparisons are often dependent on factors like sample size, marker polymorphism etc. (Aguirre-Ligouri et al., 2020; Pompanon et al., 2005). Although there are conservation studies which have suggested use of genetic indices corrected for sample size effect (Alvazrez et al., 2009; Glowatzki-Mullis et al., 2008) but their application is limited. We considered park-wise cumulative assessments of Nei’s genetic diversity (GD), allelic richness (C) and genetic erosion (Fit) indices to categorise the Indian wild rhino-bearing areas. This approach provided better quantitative option than the summary statistics-based evaluations of the parks. For example, comparative summary statistics values between MNP and DNP show contrasting results among the indices (similar Ho but higher Fis in DNP) making it difficult to ascertain the population health. However, the cumulative approach clearly categorised MNP as a genetically healthy population compared to DNP (Table xx). As the current global conservation paradigm often demands scientifically justifiable prioritization criteria (for example, conserving sites based on climatic refugia; physical distance and resistance; genetic uniqueness etc.), we demonstrated the use of genetic health indices for a species with complex history where other surrogate-based approaches e.g. environment suitability may not work (Hanson et al., 2020).

Although the recovery of Indian GOH rhino is considered as one of the successful conservation stories for large herbivores globally, managing the genetically distinct populations with almost no natural connectivity and establishing new habitats (as extant ones are reaching their respective carrying capacities) are going to be the main future challenges (Rookmaker et al., 2016; Jhala et al., 2021; Talukdar et al., 2021). Since recent Indian rhino conservation plans are focused towards translocation-driven recovery approaches (National Conservation Strategy for the Indian One-horned rhinoceros (*Rhinoceros unicornis*), Government of India, Ministry of Environment Forest and Climate Change, 2021), our simulations of future population level genetic rescue (based on available rhino demographic parameters as well as genetic health indices) provide insightful inferences for managers and policy makers. As expected, the results indicate that future maintenance of genetic diversity is dependent on genetically healthy source populations, frequency of supplementation and initial population size (for new habitats) (Jhala et al., 2021; Pacioni et al., 2020; Rummel et al., 2016; Weeks et al., 2017). Surprisingly, we found much less or no impacts of different sex ratios and duration of supplementation plan on simulated diversity, a pattern earlier shown by Jhala et al., 2021. Any opportunity to establish/maintain natural dispersal-mediated geneflow will obviously ensure better genetic rescue as observed in case of the simulation results for Orang-Kaziranga metapopulation (also shown in case of black rhinoceros by Mellya et al., 2023). However, for all remaining extant populations (except Pobitora WLS, see Figure 2 and Table S4) such natural movement events are unlikely and thus, appropriate translocation planning is the only option left to increase overall diversity and allelic richness for ensuring the species evolutionary potential.

## 5. Conclusion/Conservation Implication

By demonstrating the genetic health indices-driven population management solutions, this study argues against the standard practices of population size-based conservation efforts. Combination of sampling heterogeneity and multifaceted analyses facilitated in identifying park specific mitigation issues (Hobbs et al., 2011; Schwartz et al., 2009; Segelbacher et al., 2010), which were not simultaneously addressed in earlier landscape level studies on Indian mega fauna (De et al., 2021; Kolipakam et al., 2019; Modi et al., 2021 etc.). For example, KNP is the only park in India that may not need any genetic augmentation, and can enrich the genetic diversity of almost all the other parks (except PWLS). Similarly, Orang NP rhinos despite being moderately genetically healthy are good source population for parks with poor genetic health. In addition, Orang NP plays an important role for rhino refuges during the flooding season from Kaziranga (Talukdar 2000). As grassland encroachments and poaching pressure outside these protected areas are currently the main conservation challenges (Talukdar 2000; Talukdar et al., 2007), maintaining the natural connectivity by restoring these habitats will improve the genetic diversity of Orang NP and ensure a much higher carrying capacity for Kaziranga NP (Jhala et al., 2021). In PWLS, our simulation results suggest that any genetic rescue effort for this population is going to be largely ineffective. Therefore, major attention towards habitat management (restoration of grassland patches and reduction of grazing pressure) and human-rhino conflict mitigation is urgently required (Talukdar et al., 2007). Both genetically depauperate rhino-bearing parks of West Bengal (Ghosh et al., 2022 for mitochondrial genome and this study) require immediate population augmentation efforts (from Kaziranga NP, Orang NP and Pobitora WLS, respectively) to improve their genetic health. Finally, our data from Dudhwa NP and Manas NP show skewed genetic signatures towards one of the source populations gene-pool, thereby emphasizing the importance of multiple supplementation efforts from genetically variable source areas, followed by intensive genetic monitoring for multiple generations. Such efforts are critical for all reintroduced populations to assess the efficacy of the management plan and come up with mitigation, if required. One of the most important findings of this study is the relatively high population structure among the existing rhino populations (except Kaziranga and manas NPs), which in combination with various levels of genetic health status has led us to implement forward simulation-based scenarios to attempt ‘genetic rescue’ in this species. It is important to point out that such genetic rescue approach would help in restoring species genetic variability instead of standard management/conservation paradigm of maintaining these populations as ESU, MU etc (Ralls et al., 2018; Tallmon et al., 2004; Willi et al., 2022).

In summary, the results from this study demonstrate a useful strategy to detect early signatures of genetic erosion and potential mechanisms to address such situations in case of wild animal populations. This has been stressed highly in the Convention on Biological Diversity where assessment of genetic status (including population structure and genetic health estimates) of extant populations of endangered species was suggested for their management and future conservation. Although most of our management suggestions from this study involve genetic data driven translocation efforts to maintain viable populations for longer period of time, it is critical to remember that the concerns of grassland habitat loss are still one of the primary conservation challenges in India. Recent studies indicate alarming loss of grassland habitats from agricultural expansions, biological invasions and government mediated plantation drives leading to a decline in grassland-dependent flora and fauna (Paul et al., 2023b; Sinha et al., 2022; Vanak 2019). We hope that using rhino as a flagship species coming conservation efforts will also contribute to the protection of currently neglected grassland ecosystems (Mungi et al., 2023; Paul et al., 2023b). Finally, our study demonstrates the use of microsatellite-based population genetic methods to strategize targeted conservation efforts for a species at landscape level. If planned and implemented properly, such applications have practical use in aiding recovery plans of other endangered species in this ongoing era of Anthropocene.

## Acknowledgments

We thank the Ministry of Environment, Forest and Climate Change, Government of India for all support to implement the project. We thank the Forest Departments of Assam, West Bengal and Uttar Pradesh to give us necessary permits and field support during sampling. We are also thankful to the WWF-India team for their field and logistic support. Our thanks to Shrewshree, Dr. Rahul De, Dr. Shrutarshi Paul, Dr. Suvankar Biswas and Dr. Shiv Kumari Patel for their involvement in lab work and analysis. We thank the Director, Dean, Research Coordinator and Nodal Officer of the Wildlife Forensics and Conservation Genetics Cell for their support.

## Figure Legend

**Supplementary table 1:**
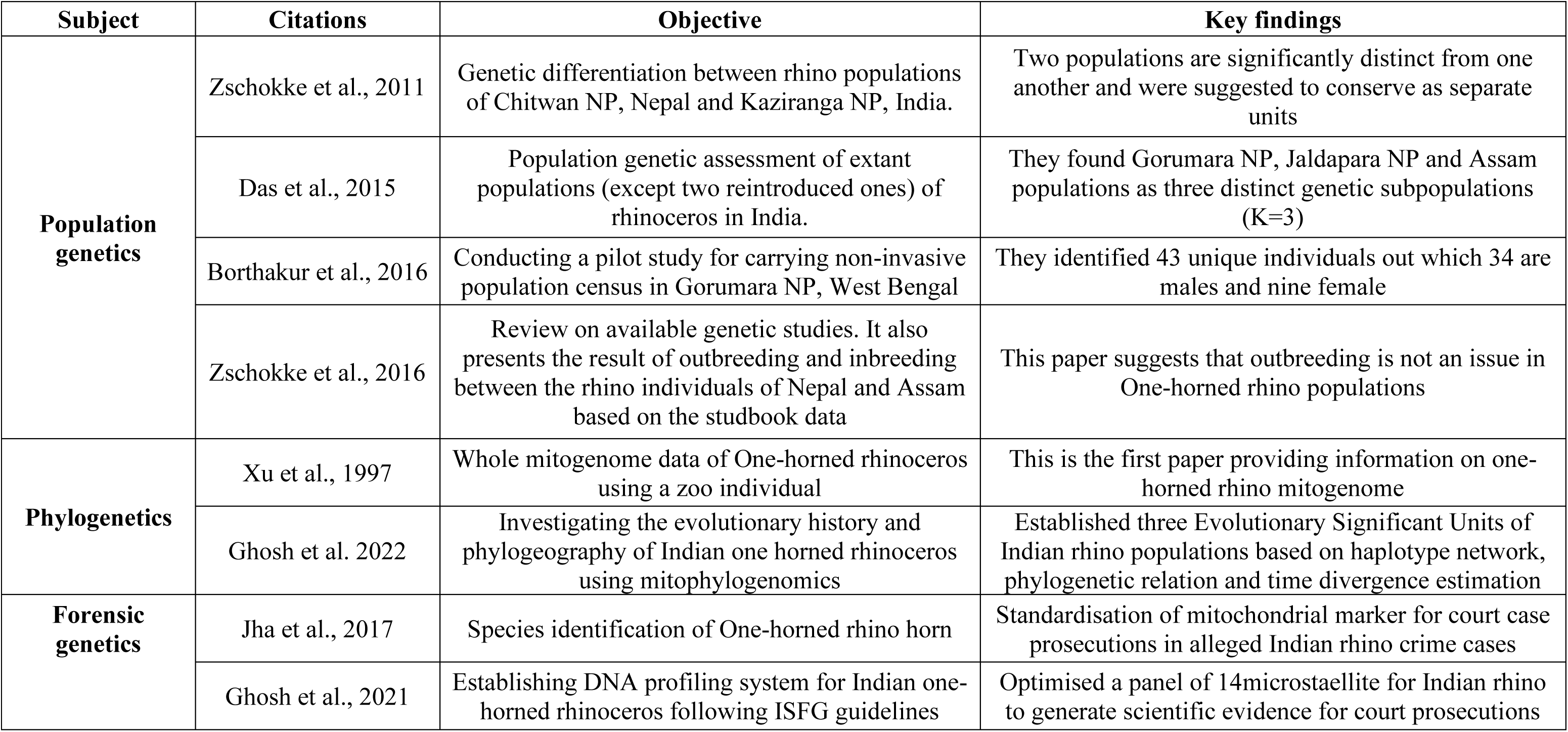
Details of the paper available on genetic studies of Greater one-horned rhinoceros.

| Subject | Citations | Objective | Key findings |
| --- | --- | --- | --- |
| **Population genetics** | Zschokke et al., 2011 | Genetic differentiation between rhino populations of Chitwan NP, Nepal and Kaziranga NP, India. | Two populations are significantly distinct from one another and were suggested to conserve as separate units |
|  | Das et al., 2015 | Population genetic assessment of extant populations (except two reintroduced ones) of rhinoceros in India. | They found Gorumara NP, Jaldapara NP and Assam populations as three distinct genetic subpopulations (K=3) |
|  | Borthakur et al., 2016 | Conducting a pilot study for carrying non-invasive population census in Gorumara NP, West Bengal | They identified 43 unique individuals out which 34 are males and nine female |
|  | Zschokke et al., 2016 | Review on available genetic studies. It also presents the result of outbreeding and inbreeding between the rhino individuals of Nepal and Assam based on the studbook data | This paper suggests that outbreeding is not an issue in One-horned rhino populations |
| **Phylogenetics** | Xu et al., 1997 | Whole mitogenome data of One-horned rhinoceros using a zoo individual | This is the first paper providing information on one-horned rhino mitogenome |
|  | Ghosh et al. 2022 | Investigating the evolutionary history and phylogeography of Indian one horned rhinoceros using mitophylogenomics | Established three Evolutionary Significant Units of Indian rhino populations based on haplotype network, phylogenetic relation and time divergence estimation |
| **Forensic genetics** | Jha et al., 2017 | Species identification of One-horned rhino horn | Standardisation of mitochondrial marker for court case prosecutions in alleged Indian rhino crime cases |
|  | Ghosh et al., 2021 | Establishing DNA profiling system for Indian one-horned rhinoceros following ISFG guidelines | Optimised a panel of 14microstaellite for Indian rhino to generate scientific evidence for court prosecutions |

**Supplementary table 2:** Brief introduction of study area in relation to the density, population size, recovery history and connectivity status of all seven rhino bearing parks of India.

| State | Park | Populati<br>on size | Density/<br>km² | Recover history | Grassland<br>area (km²) | Connectivity status | References |
| --- | --- | --- | --- | --- | --- | --- | --- |
| Uttar Pradesh | Dudhwa NP | 42 | 1.68 | Reintroduced by seven founders.<br>4 from Chitwan NP of Nepal and<br>3 from Pobitora WLS, Assam | 25 (Rhino<br>enclosure) | Isolated | Talukdar et al.,<br>2012 |
| West Bengal | Gorumara NP | 52 | 0.62 | Revived from few individuals<br>(14-19) post population crash in<br>mid 90s due to poaching | 89.05 | Isolated | Bist 1994; WB<br>report 2019 |
|  | Jaldapara NP | ~250 | 1.43 |  | 165.23 | Isolated |  |
| Assam | Manas NP | 48 | 0.28 | Reintroduced from 2010-2020<br>through multiple<br>supplementations from Kaziranga<br>NP (12), Pobitora WLS (10) and<br>CWRC (13) | 174 | Isolated | Talukdar et al.,<br>2021; Sinha et al.,<br>2022 |
|  | Pobitora WLS | 101 | 6.25 | 54 individuals recorded in 1987<br>after Wildlife sanctuary<br>declaration. Forest department<br>records shows presence of 14 odd<br>rhinos during 1961-1962 | 16 | Isolated but there are<br>reports of rhinos<br>straying out to Orang<br>NP during monsoon | Talukdar 2000;<br>Talukdar et al.,<br>2007 and 2021 |
|  | Orang NP | 100 | 1.26 | Localised by animals from<br>Kaziranga NP post Laeishmania<br>calamity in 1900 | 79 | Connected via patches<br>of grasslands along<br>Brahmaputra<br>floodplains. Since<br>Orang is highland,<br>many times rhinos of<br>KNP are fled here<br>post monsoon |  |
|  | Kaziranga NP | ~2500 | 5.53 | Revived from few hundred<br>individuals post population crash<br>in mid 90s due to poaching<br>pressures | 432 |  |  |

**Supplementary Table 3:** Summary of Bayesian clustering analysis of Indian rhino populations and their respective estimates of inferred genetic clusters.

| Bayesian IBC analysis | Models | Inferred K | Genetic sub-populations |  |
| --- | --- | --- | --- | --- |
|  |  |  | Unique populations | Mixed populations |
| STRUCTURE | Non spatial, Admixture | 4 | GNP, JNP, KNP and PWLS | DNP=PWLS; MNP=KNP; ONP= mix of KNP and PWLS |
|  | Spatial, Admixture | 5 | DNP,GNP, JNP, KNP and PWLS | MNP=KNP; ONP= mix of KNP and PWLS |
| TESS | Spatially explicit, Admixture | 5 |  |  |
| GENELAND | Spatially explicit, Non-admixture | 7 | DNP,GNP, JNP, MNP,PWLS, ONP, KNP | NO |

**Supplementary table 3:**
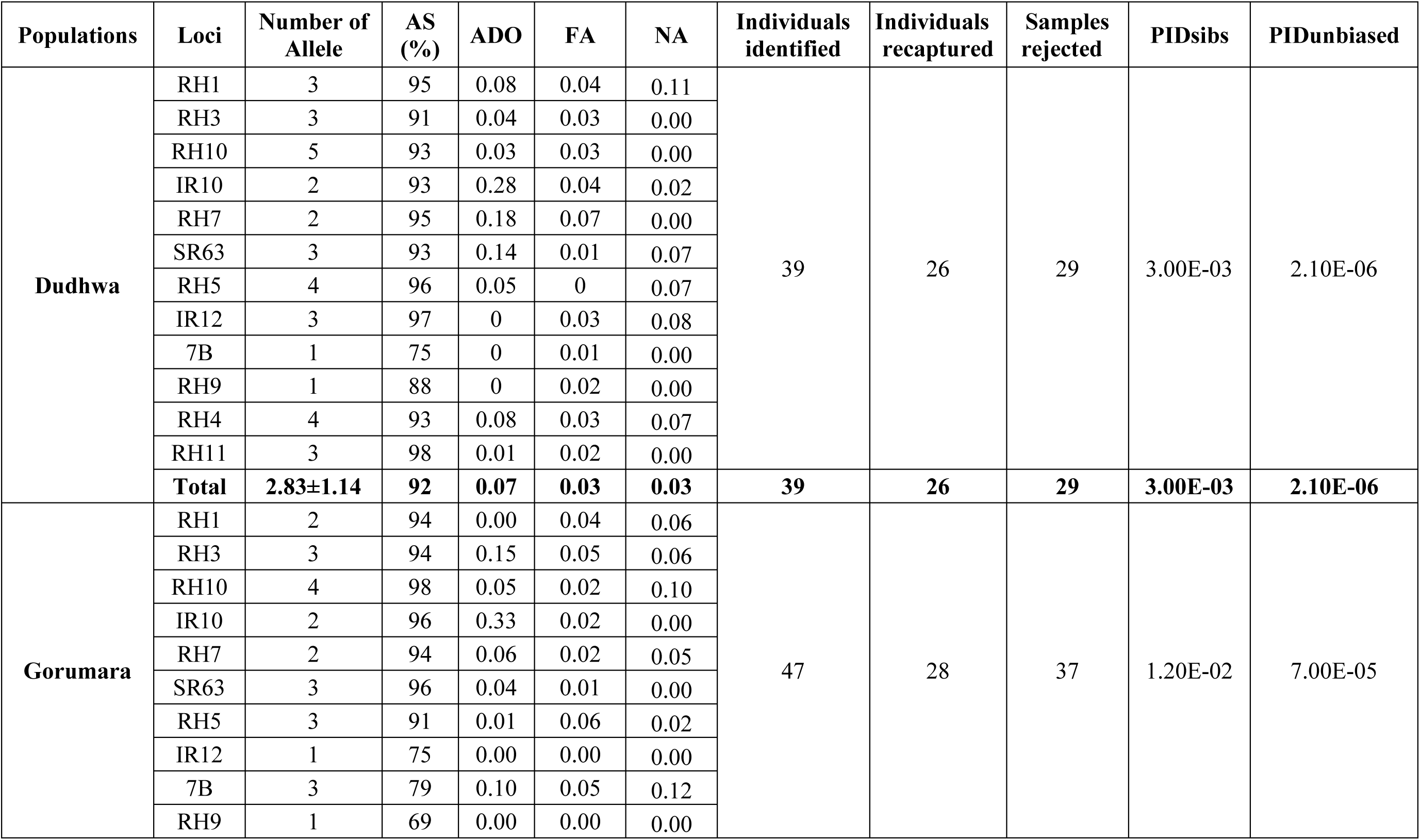
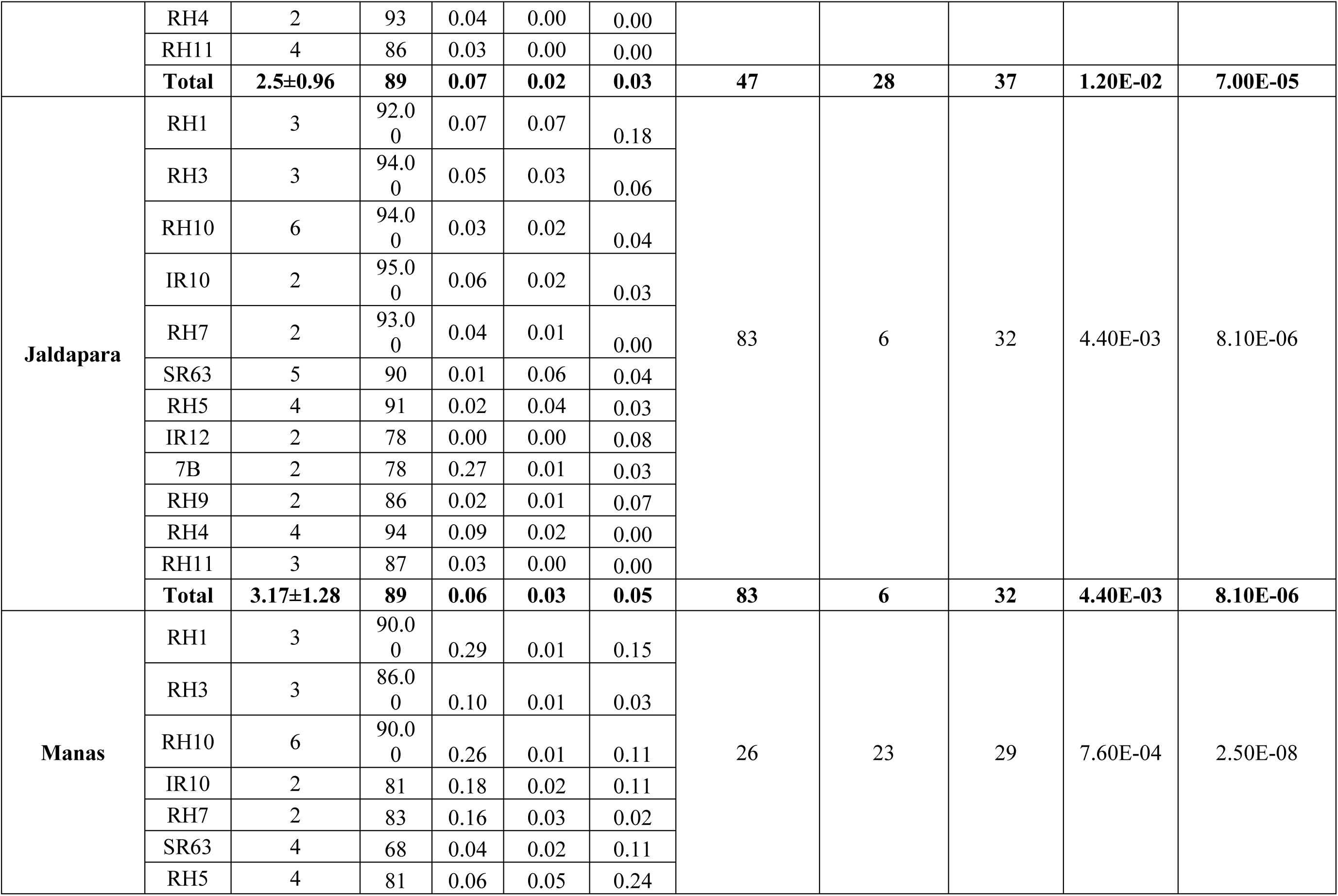
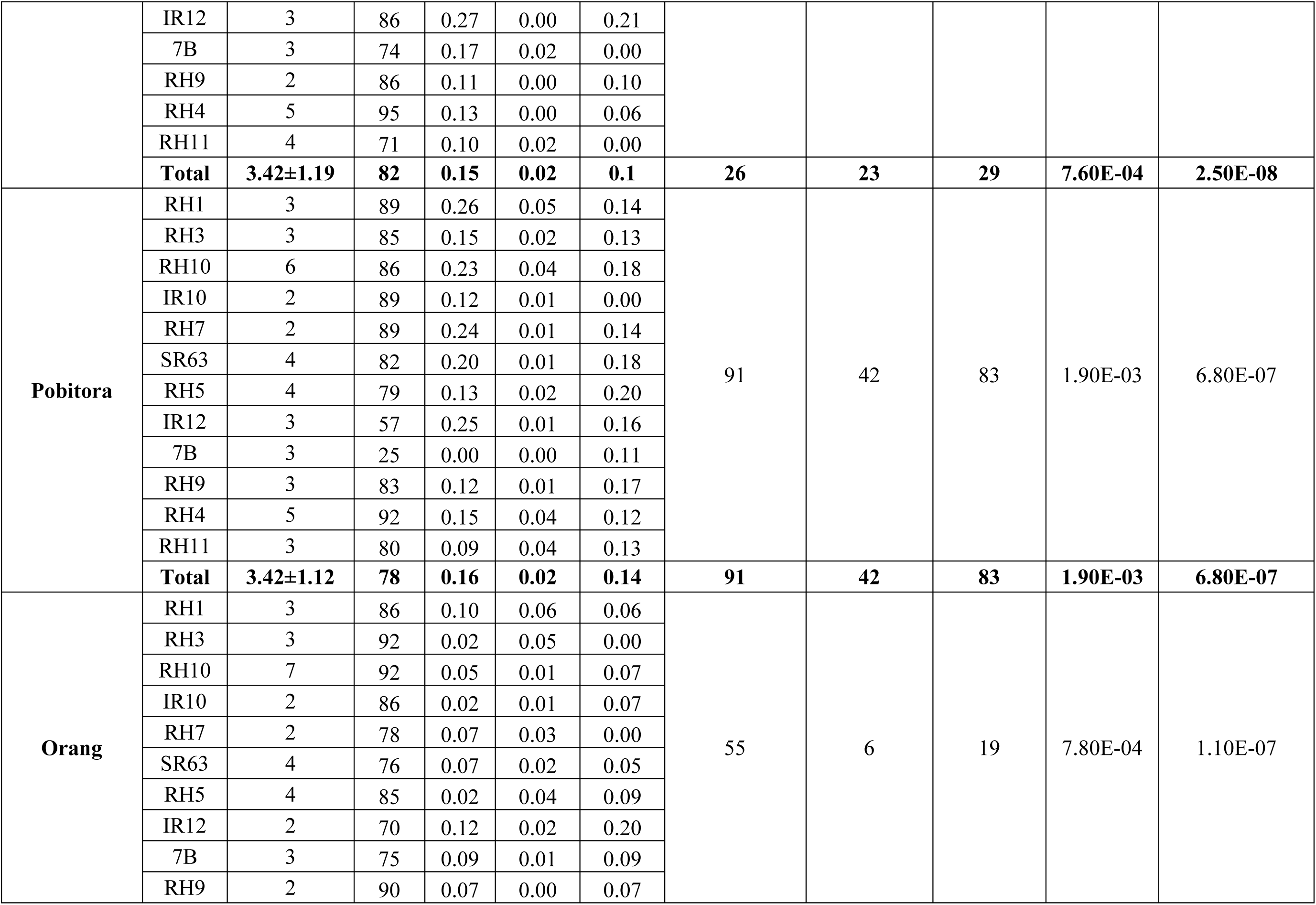
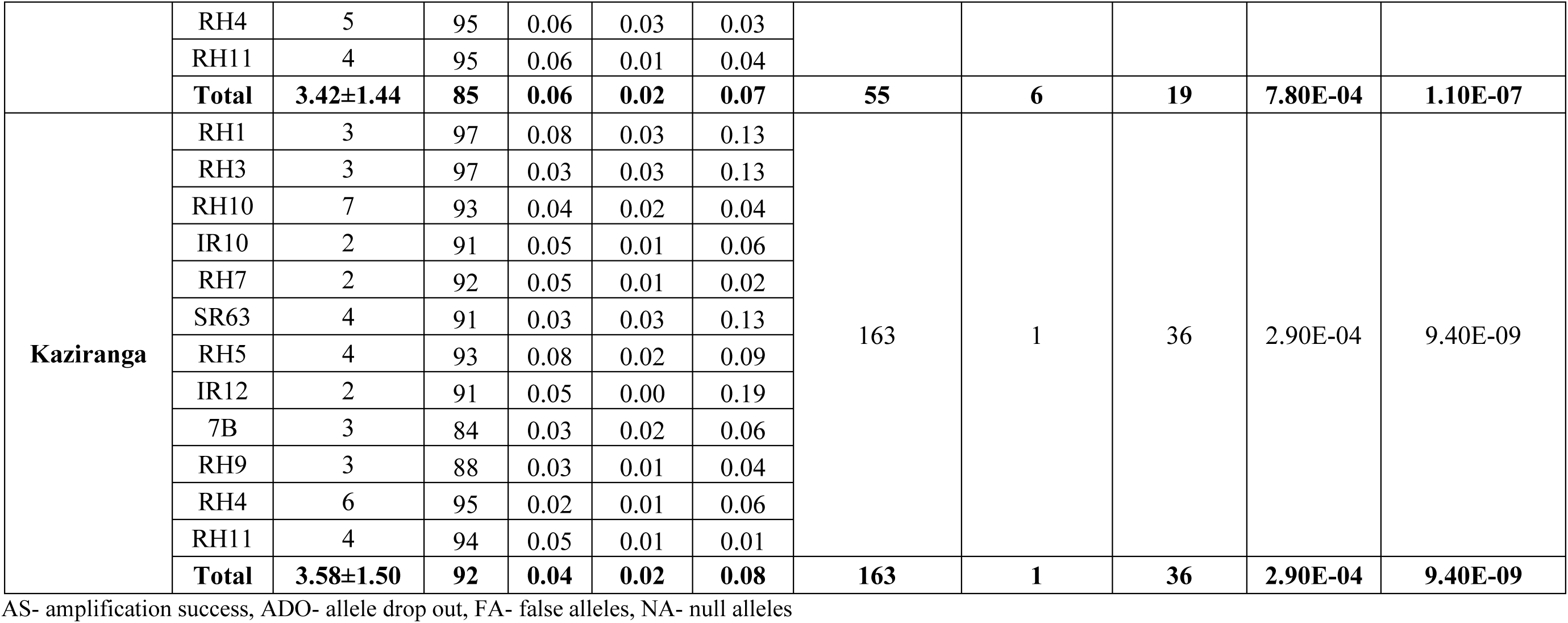
Details of loci wise summary statistics across all Indian rhino populations.

**Supplementary tab.**
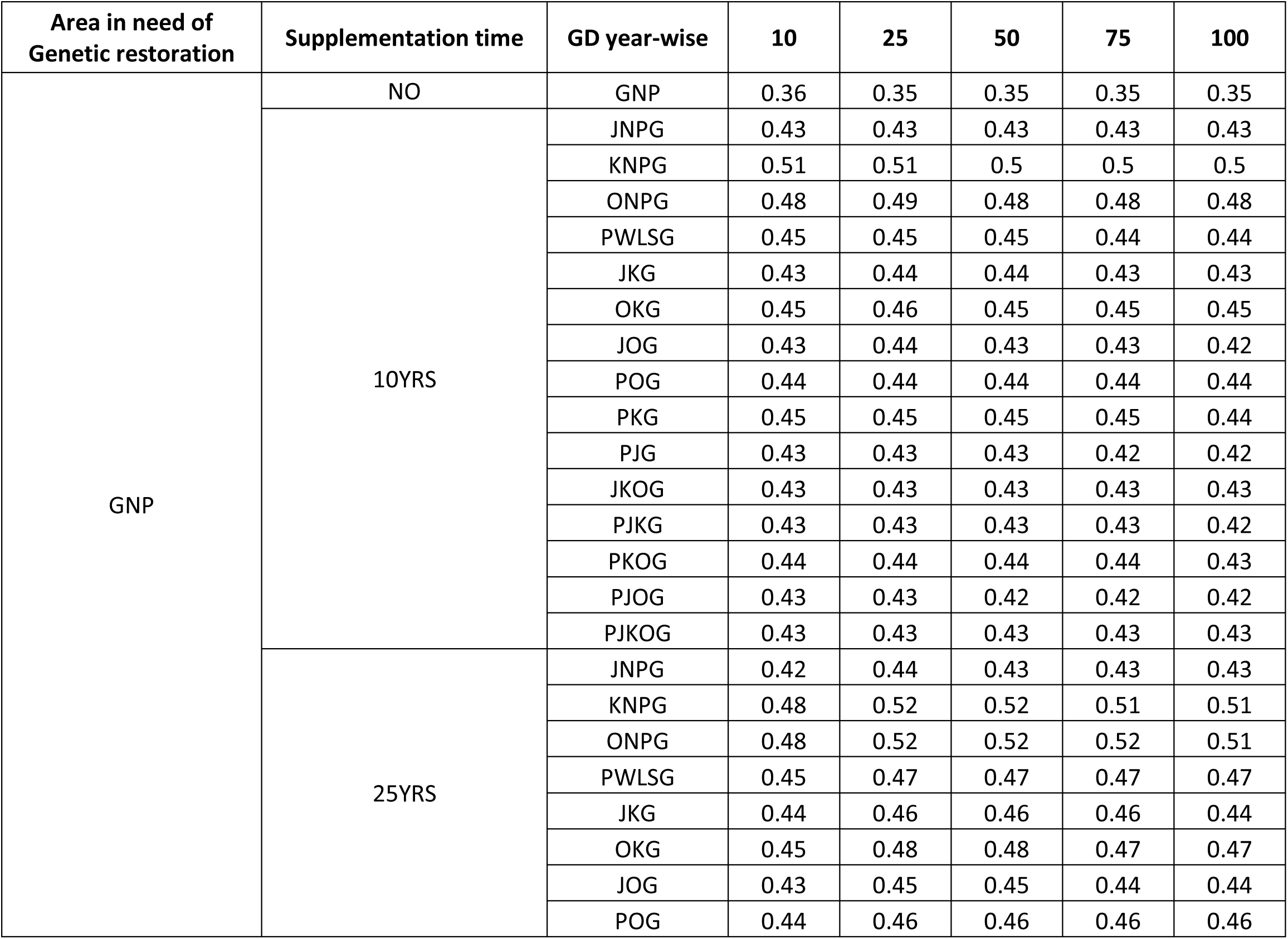
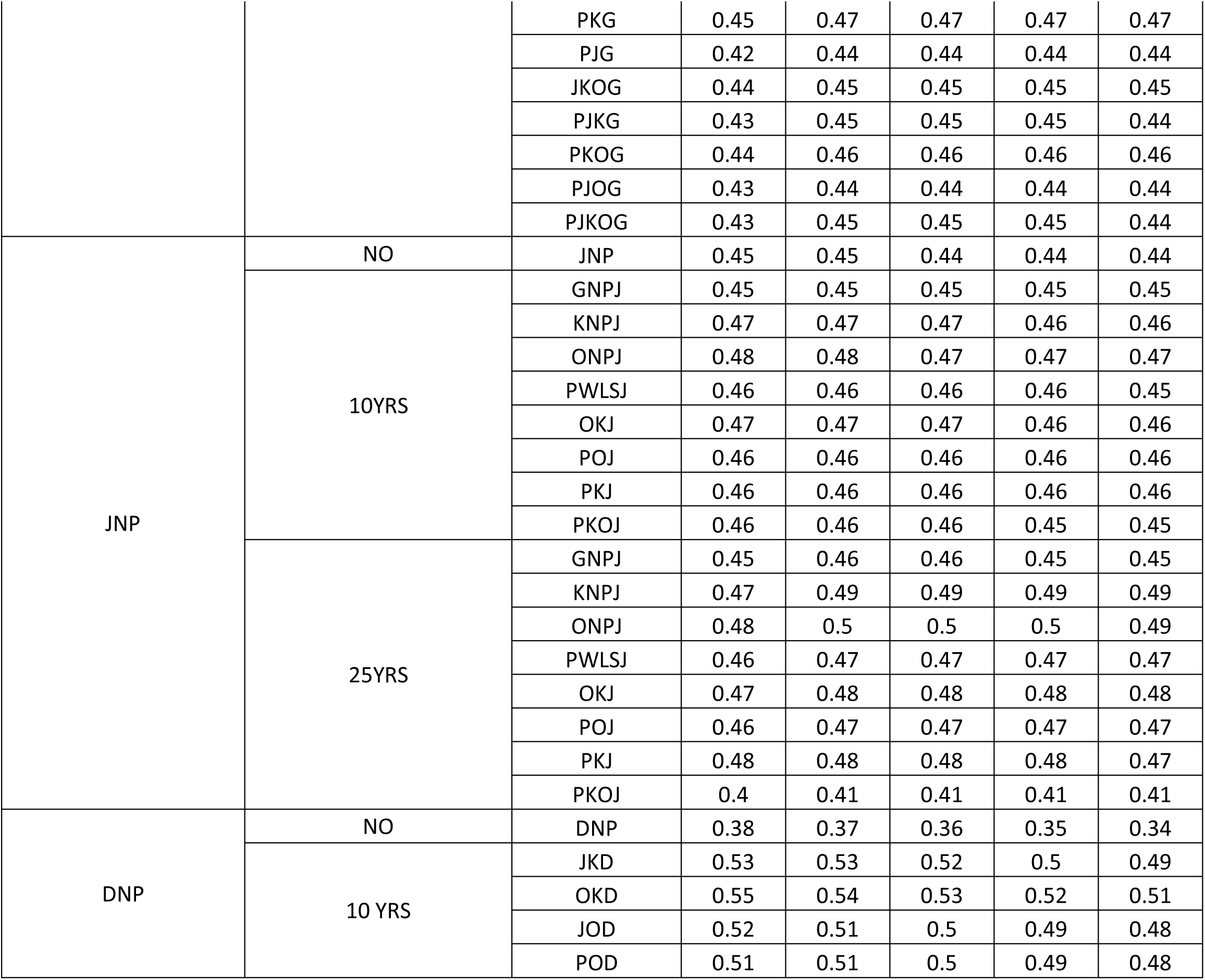
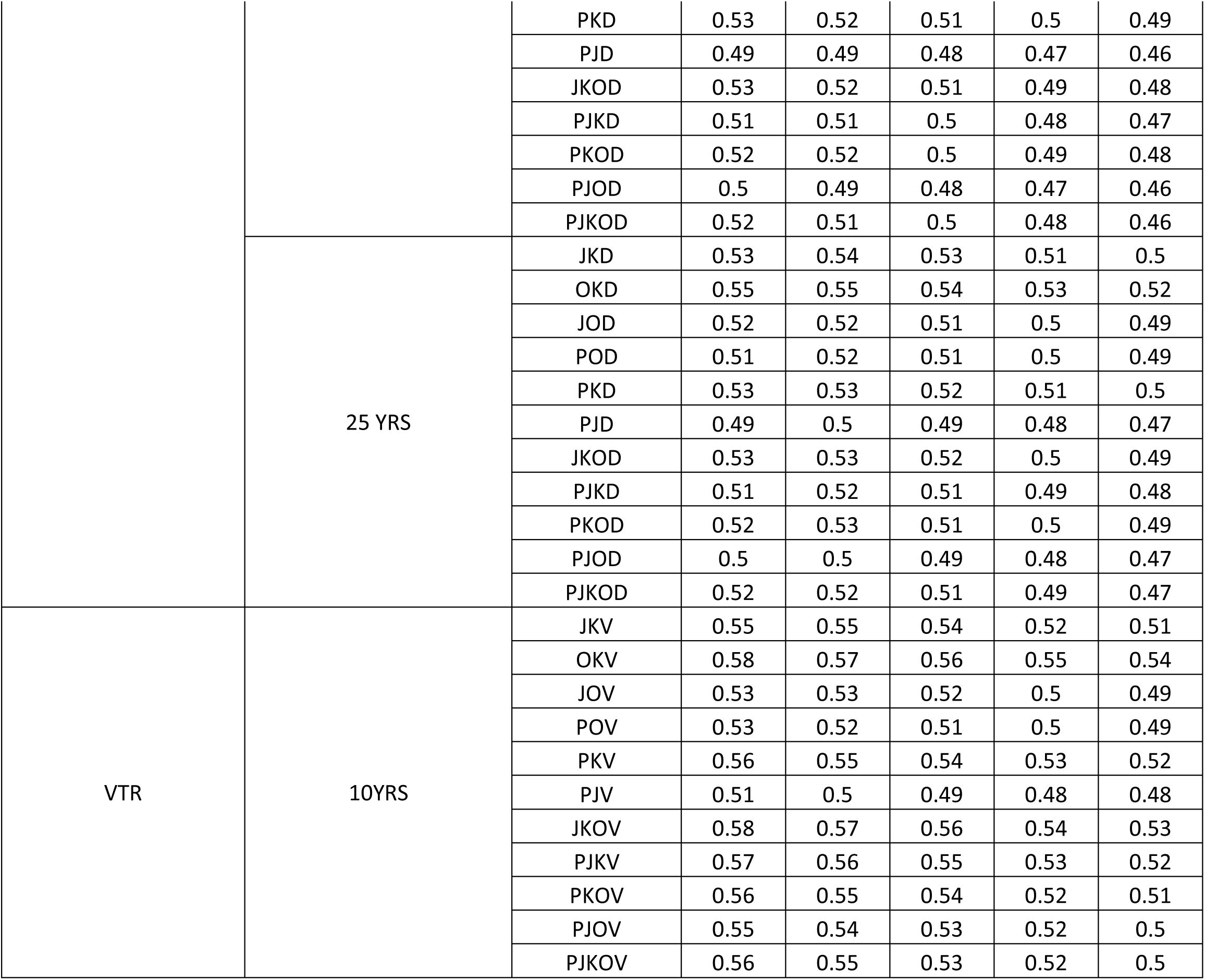
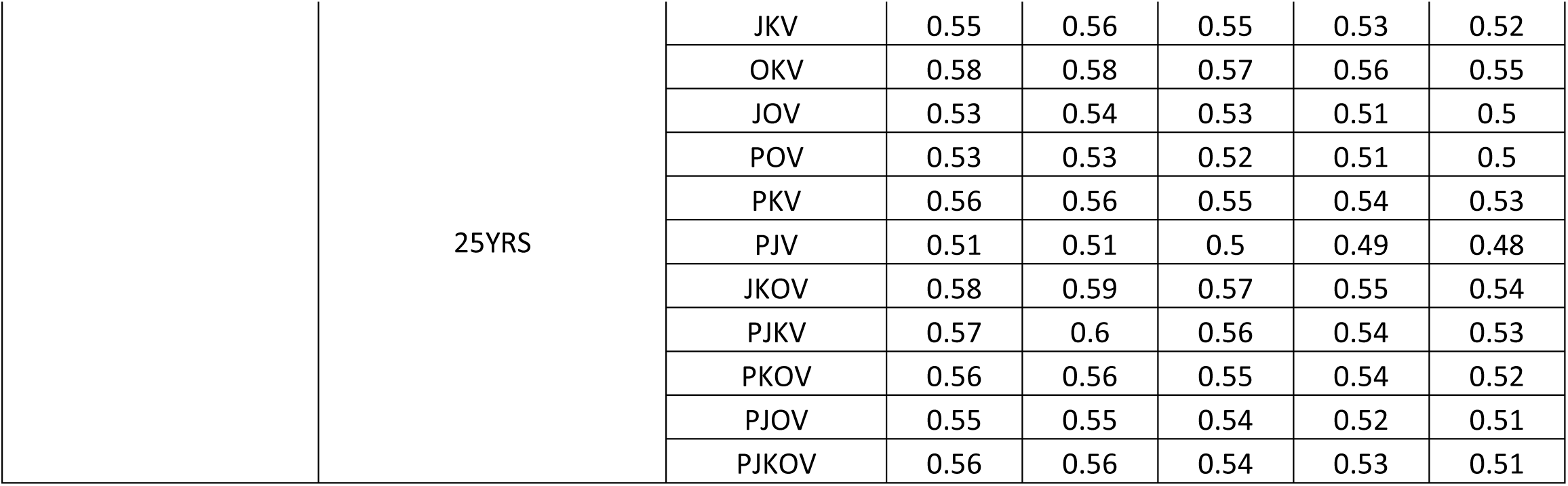
Table 4: Details of forward simulations for managed translocation plans in new potential habitats and extant populations with poor health.

**Supplementary figure 1:**
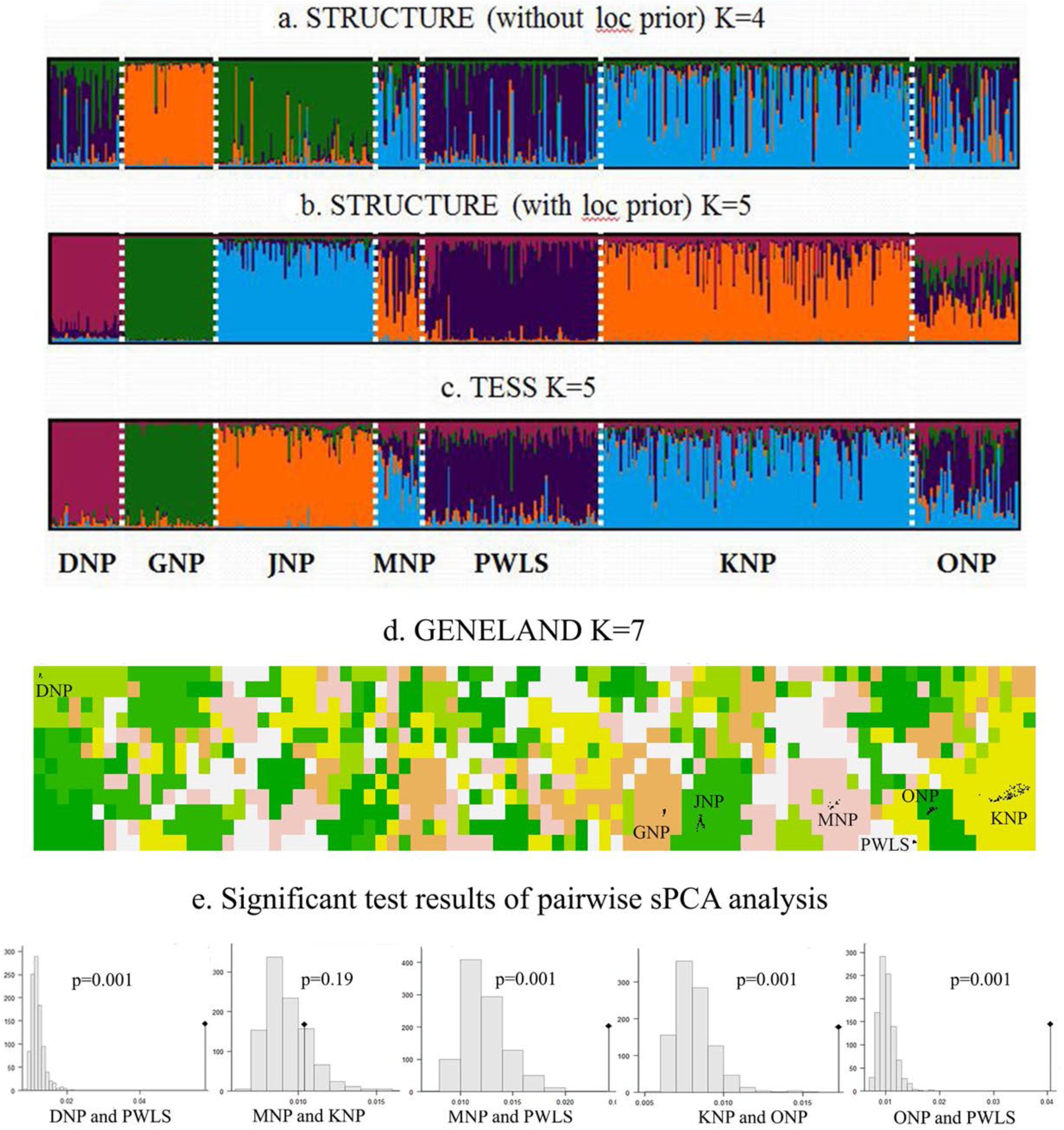
Results of integrated multiple clustering approach to identify the genetic subpopulation at landscape level. In STRUCTURE optimal K is decided using model choice statistics L(K) which concluded in (a) K=4 for non-loc prior run and (b) K=5 with loc-prior. For TESS Kmax is finalised using the DIC value resulting in K=5 (c). GENELAND shows Indian rhino populations are genetically clustered into 7 sub-populations represented by population membership graph (d). The optimal K was decided based on the highest mean posterior density among each candidate K. Finally sPCA analysis was done to identify the misinterpreted subpopulation by IBC approach (e). The result shows that Manas is not a different genetic sub-population (p=0.19) and thus GENELAND was over estimating it. On the other hand, Orang is a genetic sub-population which could not be identified using admixture-based models (STRUCTURE and TESS) thus concluding into six genetic sub-populations in Indian rhinos.

**Supplementary figure 2:**
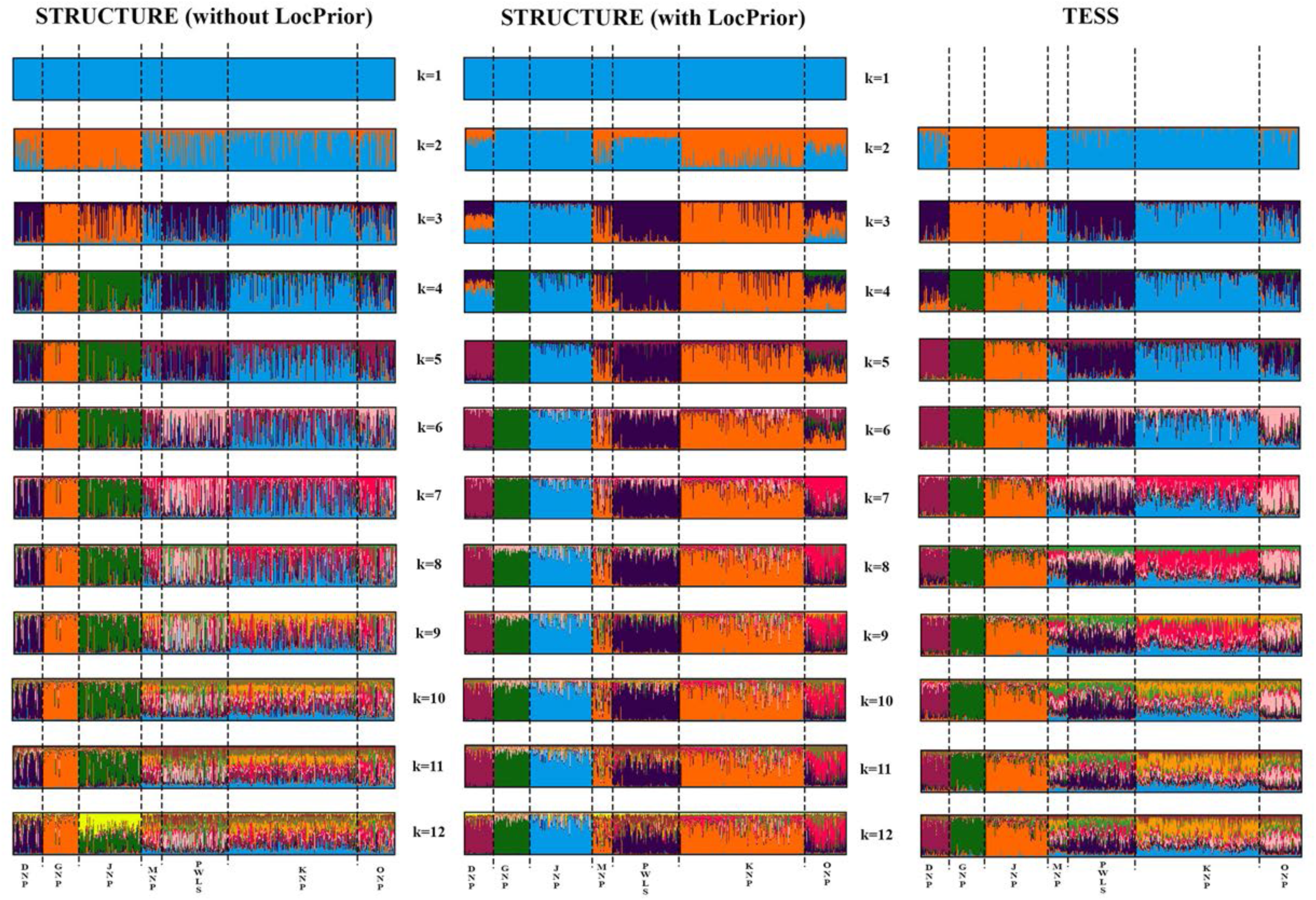
All the bar-plots of STRUCTURE (with and without loc prior) and TESS run for K= 1-12.

